# *Nf1* deletion results in depletion of the *Lhx6* transcription factor and a specific loss of parvalbumin+ cortical interneurons

**DOI:** 10.1101/746214

**Authors:** Kartik Angara, Emily Ling-Lin Pai, Stephanie M Bilinovich, April M Stafford, Julie T Nguyen, Anirban Paul, John L Rubenstein, Daniel Vogt

## Abstract

Neurofibromatosis-1 (NF-1) is a monogenic disorder caused by mutations in the *NF1* gene, which encodes the protein, Neurofibromin, an inhibitor of Ras GTPase activity. While NF-1 is often characterized by café-au-lait skin spots and benign tumors, the mechanisms underlying cognitive changes in NF-1 are poorly understood. Cortical GABAergic interneurons (CINs) are implicated in NF-1 pathology but cellular and molecular changes to CINs are poorly understood. We deleted *Nf1* from the medial ganglionic eminence (MGE), which gives rise to both oligodendrocytes and CINs that express somatostatin and parvalbumin. Loss of *Nf1* led to a persistence of immature oligodendrocytes that prevented later born oligodendrocytes from occupying the cortex. Moreover, PV+ CINs were uniquely lost, without changes in SST+ CINs. We discovered that loss of *Nf1* results in a graded decrease in *Lhx6* expression, the transcription factor necessary to establish SST+ and PV+ CINs, revealing a mechanism whereby *Nf1* regulates a critical CIN developmental milestone.

## Introduction

Neurofibromatosis-1 (NF-1) is caused by mutations in the gene, *NF1*, which encodes the Neurofibromin protein (Mattocks et al., 2004). Neurofibromin contains a centrally located GTPase activating protein (GAP) domain, which inhibits RAS GTPase activity (Klose et al., 1998), but also contains many other regions for unique protein-protein interactions that could underlie different cellular functions (Ratner and Miller, 2015). NF-1 is one of many RAS-opathies (Rauen et al., 2010), and is often diagnosed by the appearance of skin discoloration (café-au-lait spots) and benign tumors, however, there are also cognitive changes, including a high incidence of autism spectrum disorder (ASD) and learning disabilities (Garg et al., 2013; Morris et al., 2016; Vogel et al., 2017). While the link between RAS-opathies and ASD has been documented (Vithayathil et al., 2018), the cellular and molecular mechanisms underlying how these cognitive changes arise are poorly understood.

In animal models, loss of *Nf1* leads to altered myelination, increased oligodendrocyte precursors in the spinal cord and preferential gliogenesis at the expense of neurogenesis in the brain (Bennett et al., 2003; Lee et al., 2010; Mayes et al., 2013; Wang et al., 2012a). Moreover, a pan-GABAergic deletion of *Nf1* in mice was shown to underlie cognitive performance and altered excitatory/inhibitory balance (Costa et al., 2002; Cui et al., 2008). Overall, these studies suggest changes in glia and/or GABAergic neurons as potential modulators of *Nf1*’s function in the brain. However, a detailed molecular and cellular study of both these cell types after loss of *Nf1* is lacking.

The median ganglionic eminence is a unique telencephalic progenitor domain that gives rise to multiple cell types, including the majority of GABAergic cortical interneurons (CINs) (Wonders and Anderson, 2006) and the first wave of oligodendrocytes to populate the brain (Kessaris et al., 2006). MGE-derived oligodendrocytes die off after birth and later waves of oligodendrocytes that are derived from *Gsh2* and *Emx1* lineages are the dominant population in the adult cortex (Kessaris et al., 2006). Moreover, MGE-lineage oligodendrocytes are not well understood and their role during development and their contribution to cortical development are unclear. MGE-derived CINs primarily express the molecular markers somatostatin (SST) or parvalbumin (PV), which are determined by the transcription factor, *Lhx6*, after the MGE is regionally patterned by *Nkx2.1* (Liodis et al., 2007; Sussel et al., 1999; Zhao et al., 2008). *Lhx6* is a cardinal cell fate determination factor expressed in the MGE and when deleted, SST+ and PV+ CINs do not acquire their mature cell fates and a subset assume properties of distinct caudal ganglionic eminence (CGE)-derived CINs (Liodis et al., 2007; Silberberg et al., 2016; Vogt et al., 2014; Zhao et al., 2008). Of note, the CGE gives rise to other classes of CINs, including the vasoactive intestinal peptide (VIP) expressing and the REELIN+/SST-groups (Miyoshi et al., 2010). While recent studies have examined the role of transcription factors that influence MGE and CGE CIN development (Hu et al., 2017a; Mayer et al., 2018; Pai et al., 2019), few reports have assessed how cellular signaling proteins regulate these processes (Malik et al., 2019; McKenzie et al., 2019). After their initial programming in the MGE, CINs tangentially migrate into and laminate the neocortex. As they mature, they become diversified into further subclasses with distinct molecular, anatomical and physiological properties (Hu et al., 2017b; Kessaris et al., 2014). These diverse properties contribute unique roles to brain function but whether distinct CIN groups are altered in NF-1 is unknown.

Since both oligodendrocytes and GABAergic CINs have been implicated in NF-1, the MGE provides a unique opportunity to probe both cell populations after *Nf1* deletion. Herein, we show loss of *Nf1* in mouse MGE-lineages results in a persistence of immature oligodendrocytes in the adult neocortex, preventing later born oligodendrocytes from occupying the same region. This was accompanied by loss of over half of the CINs that express PV, without any changes in the numbers of CINs that express SST, suggesting a specific effect on this GABAergic CIN cell population. We also found that *Lhx6* was depleted in a dose-dependent manner when *Nf1* was deleted from MGE cells, providing a new mechanism for how *Nf1* dysfunction could alter the normal development of CINs. This provides insights into both glial and CIN molecular and cellular alterations that could occur in NF-1 and suggest a novel mechanism linking *Nf1* loss to alterations in a cardinal CIN developmental program that may underlie some cognitive changes in NF-1.

## Results

### Conditional loss of *Nf1* from the MGE results in a persistence of immature MGE-lineage oligodendrocytes

To assess *Nf1*’s role in MGE-lineages, we crossed *Nf1^Floxed^* mice (Zhu et al., 2001) with *Nkx2.1-Cre*, which begins to express ~embryonic day (E) 9.5 in progenitors of the MGE (Xu et al., 2008a); these crosses also included the *Ai14* allele, which expresses tdTomato after cre-recombination (Madisen et al., 2010). The conditional heterozygous (cHet) and knockout (cKO) mice were born at Mendelian ratios, and survived into adulthood. To validate this conditional deletion, we used fluorescent in situ hybridization (FISH) to detect *Nf1* in a subset of MGE-lineage CINs, i.e. SST+, in the somatosensory cortex at postnatal day (P)30. *Nf1* transcript was depleted from these cells in cKOs (Figure S1A).

Next, we collected *Nf1* wild type (WT), conditional Heterozygous (cHet) and homozygous (cKO) brains at P30 and assessed *Nkx2.1-Cre*-lineage cells in the neocortex. While WT and cHet brains looked equivalent, cKO brains had an increase in *Nkx2.1-Cre*-lineage cells (tdTomato+) with small cell bodies (Figures 1A-1F). Previous studies have indicated that glial numbers can increase after *Nf1* deletion (Bennett et al., 2003; Lee et al., 2010; Wang et al., 2012a). Since the early wave of oligodendrocytes are derived from the MGE (Kessaris et al., 2006), we explored whether oligodendrocytes may represent this population. Thus, we first probed for OLIG2, which is a general marker of oligodendrocytes. While we did not find any differences in the total number of Olig2+ cells (Figure 1G), there was ~200-fold increase in the number of tdTomato+ cells co-labeled for Olig2 in the cKOs (Figures 1A-1C and 1F, p < 0.0001 compared to both WT and cHet).

**Figure 1.**
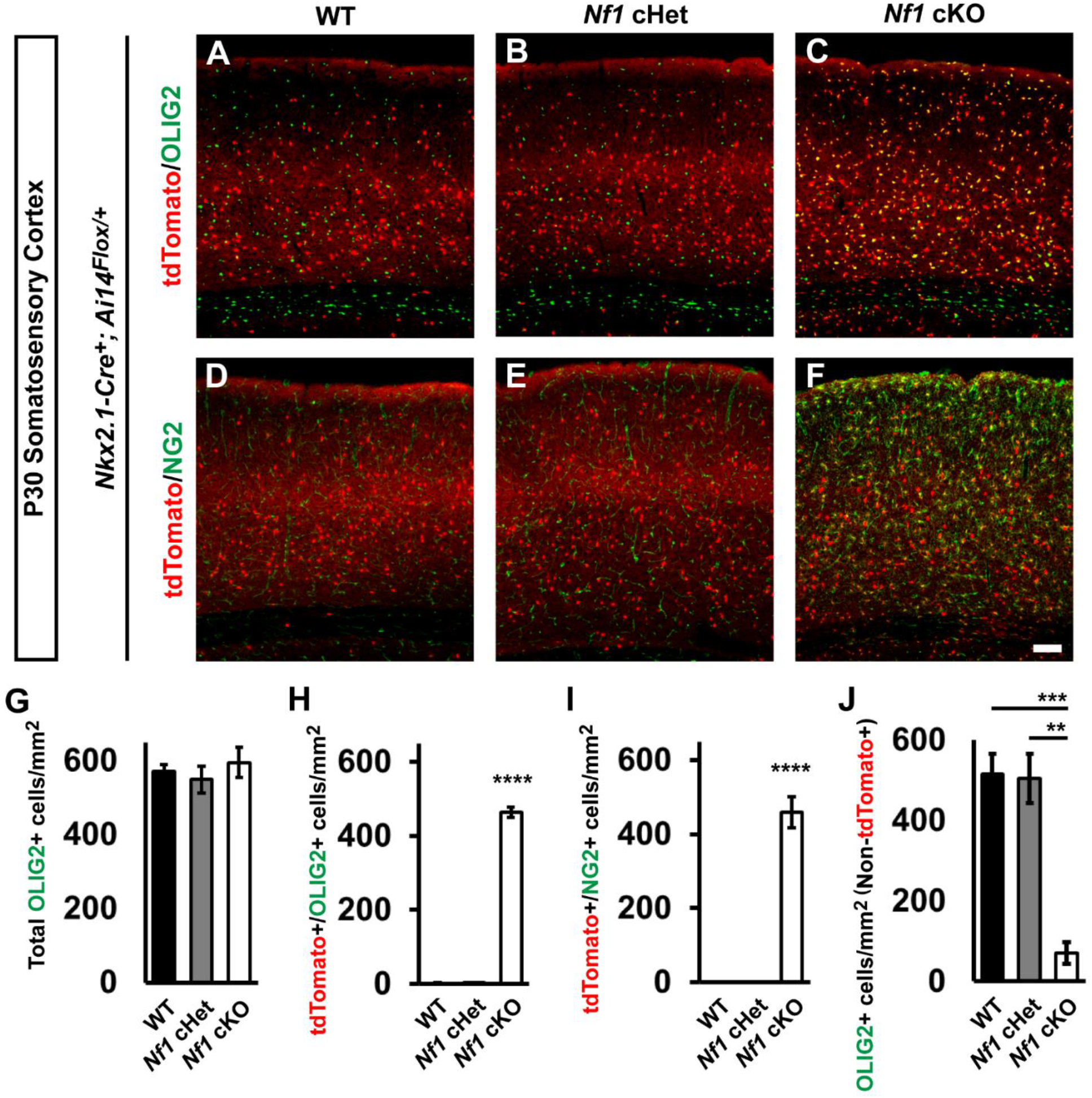
Persistence of immature *Nkx2.1-Cre*-lineage and depletion of later born oligodendrocytes in *Nf1* cKOs. Immunofluorescent images from the somatosensory cortex of P30 *Nkx2.1-Cre^+^; Ai14^Flox/^*^+^ mice that were either WT (**A** and **D**), *Nf1^Flox/+^* (cHet) (**B** and **E**), or *Nf1^Flox/Flox^* (cKO) (**C** and **F**). Images show *Nkx2.1-Cre*-lineages (tdTomato+) co-labeled for the oligodendrocyte markers OLIG2 (all oligodendrocytes) or NG2 (oligodendrocyte progenitor cells, OPCs). (**G**) Quantification of the cell density of total OLIG2+ cells in the somatosensory cortex at P30. (**H**) Quantification of double-labeled tdTomato^+^/OLIG2^+^ cells in the somatosensory cortex at P30. (**I**) Quantification of the cell density of double-labeled tdTomato^+^/NG2^+^ cells in the somatosensory cortex at P30. (**J**) Quantification of OLIG2+ cells that were not tdTomato+ in the somatosensory cortex at P30. Data are presented as the mean ± SEM. All groups, n = 4. ** p < 0.01, *** p < 0.001, **** p < 0.0001. Scale bar in (**F**) = 100 μm.

By P30, MGE-lineage oligodendrocytes should not be present in the somatosensory cortex, as they die off after birth (Kessaris et al., 2006). Since it is unknown how mature these MGE-derived oligodendrocytes are, we probed for NG2, a marker of oligodendrocytes precursor cells (OPCs) that are still immature. Interestingly, many of the cKO MGE-lineage oligodendrocytes were NG2+, suggesting they were still OPCs; there were no NG2+ cells in the WTs or cHets (Figures 2D-2F and 2I, p < 0.0001 compared to both WT and cHet). Finally, we asked what effect these immature oligodendrocytes from the MGE had on the later born waves of oligodendrocytes in the somatosensory cortex. Thus, we counted the Olig2+ cells that lacked tdTomato expression and found ~84% reduction in oligodendrocytes that were not tdTomato+ in cKOs (Figure 2J, WT, p = 0.0009 and cHet, p = 0.001). Thus, the persistence of early born MGE-lineage oligodendrocytes prevented later born waves from occupying the cortex.

**Figure 2.**
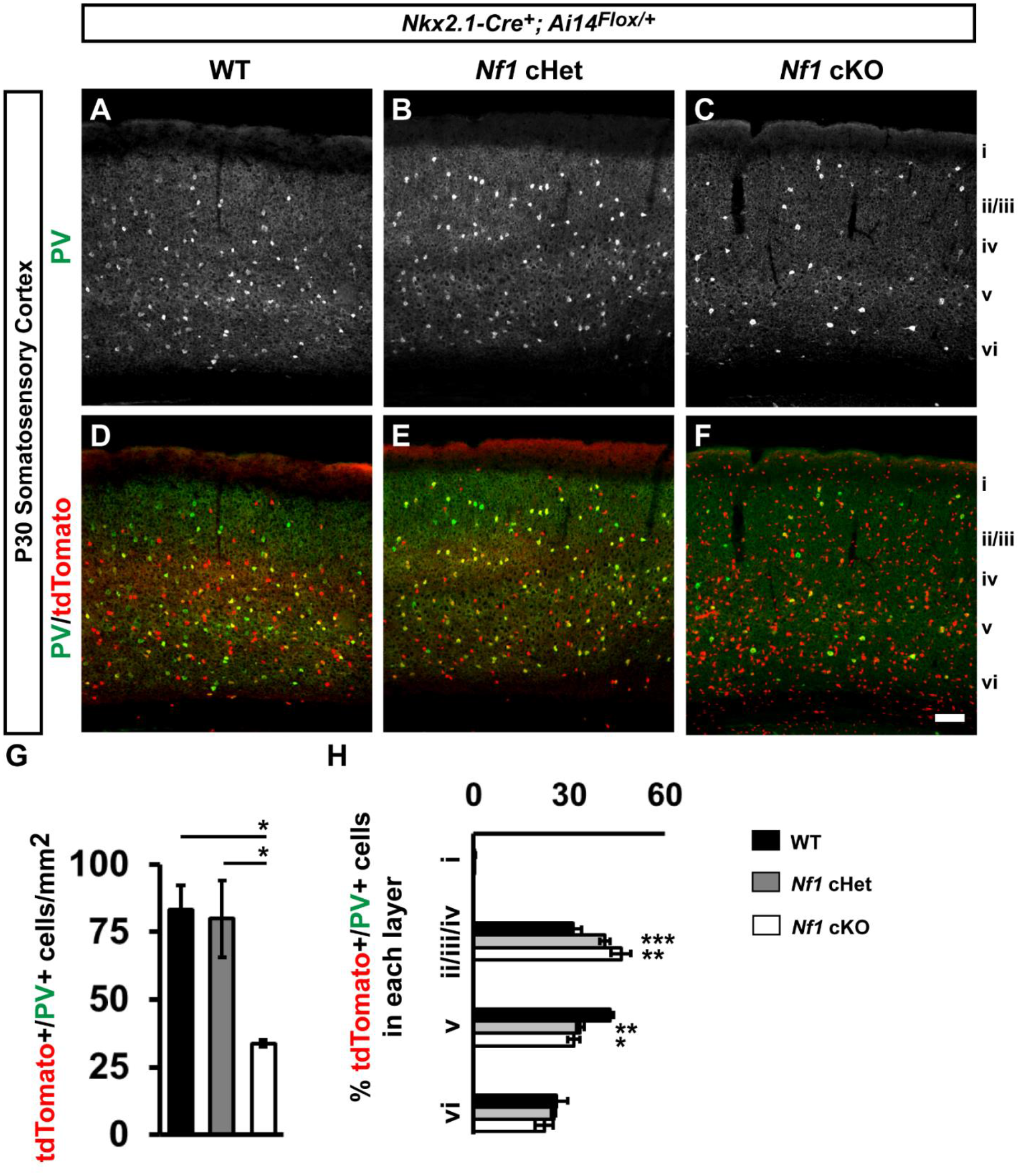
Unique loss of PV+ cortical interneurons in *Nf1* cKOs. Immunofluorescent images from the somatosensory cortex of P30 *Nkx2.1-Cre^+^; Ai14^Flox/^*^+^ mice that were either WT (**A, A’, B, B’**), *Nf1^Flox/+^* (cHet) (**C, C’, D, D’**), or *Nf1^Flox/Flox^* (cKO) (**E, E’, F, F’**). Images show *Nkx2.1-Cre*-lineages (tdTomato+) co-labeled for the MGE-derived cortical interneuron markers somatostatin (SST), or parvalbumin (PV). Roman numerals to the right of the images denote cortical lamina. Quantification of the cell density of double-labeled tdTomato^+^/SST^+^ cells (**G**) or tdTomato^+^/PV^+^ cells (**H**) in the somatosensory cortex at P30. Quantification of the cortical laminar distribution of double-labeled tdTomato^+^/SST^+^ cells (**I**) or tdTomato^+^/PV^+^ cells (**J**). Data are presented as the mean ± SEM. All groups, n = 4. * p < 0.05, ** p < 0.01, *** p < 0.001. Scale bar in (**F’**) = 100 μm.

### PV+ CINs are specifically reduced in *Nf1* cKO somatosensory cortex

We next asked if *Nf1* loss altered the development of MGE-derived CINs. To this end, we first wanted to establish whether *Nf1* was expressed in different CIN groups. Thus, we examined single-cell levels of *Nf1* transcripts from ~P30 CINs in the cortex (Paul et al., 2017). *Nf1* transcripts were found in all CIN cell types examined, including both MGE and CGE derived groups (Figure S1B). This suggested that *Nf1* might have a role in each group of CIN; *Nkx2.1-Cre* will delete *Nf1* in both SST and PV CINs, which represent the majority of CINs.

We probed for SST and PV expression in the somatosensory cortices of *Nf1* WTs, cHets and cKOs at P30 (Figures 2A-2F). Interestingly, while there was no difference between WT and cHet PV+ CINs, cKOs had ~60% decrease in PV+ CINs (Figure 2G, vs. WT, p = 0.03 and cHet, p = 0.04). In addition, there was a greater proportion of PV+ CINs in cortical layers ii/iii/iv and less in layer v in both cHets and cKOs (Figure 2H, layers ii/iii/iv WT vs. cHet p = 0.0007 and WT vs. cKO p = 0.002, layer v WT vs. cHet p = 0.005 and WT vs. cKO p = 0.01). In contrast, we did not detect any changes in SST cell density or lamination between genotypes (Figure S2). This demonstrates that even though both SST and PV CINs express *Nf1*, the PV group is particularly sensitive to *Nf1* loss.

### MGE-lineage oligodendrocyte and PV+ CIN phenotypes are also observed in the hippocampus and striatum

We next assessed for MGE-lineages in the hippocampus at the same age. Similar to somatosensory cortex, hippocampal MGE-derived OLIG2+ cells were increased ~17-fold in *Nf1* cKOs Figures (S3A, S3D, S3G and S3J, p < 0.0001). Whereas SST+ interneurons were not significantly different (Figures S3B, S3E, S3H and S3K), the PV+ interneurons were reduced in *Nf1* cKOs by 42% (Figures S3C, S3F, S3I and S2L, p = 0.01).

We also examined MGE-lineage cells in the striatum and found that MGE-derived lineages had ~45-fold increase in tdTomato+/OLIG2+ cells in the *Nf1* cKOs (Figures S4A, S4E, S4I and S4M, p = 0.0004 compared to both WT and cHet). Since cholinergic interneurons are also derived from the MGE, we assessed the number of choline acetyltransferase (ChAT)+ cells in striatum but found no difference between groups (Figures S4B, S4F, S4J and S4N). In addition, the numbers of SST+ striatal interneurons were unchanged (Figures S4C, S4G, S4K and S4O). However, consistent with other brain regions, we found that *Nf1* cKOs had ~32% decrease in striatal PV+ interneurons (Figures S4D, S4H, S4L and S4P, p = 0.04). These data show that loss of *Nf1* affects the hippocampus and striatum in a similar manner to the somatosensory cortex.

### No change in PV+ CINs when *Nf1* is deleted from post-mitotic *PV-Cre*-lineage CINs

To determine if *Nf1* was necessary in post-mitotic PV+ CINs, we crossed our *Nf1^Floxed^; Ai14* mice with a *PV-Cre* line (Hippenmeyer et al., 2005). We first calculated the cell density of *PV-Cre*- lineage CINs (tdTomato+) in the somatosensory cortex at P30 and found no differences between genotypes (Figures 3A, 3D, 3G and 3J). We also co-labeled these cells with PV and found no difference in the number of CINs that expressed PV (Figures 3A-3I and 3K). We also co-labeled for SST, which should not be expressed in *PV-Cre*-lineages and found no expression of SST in any group (data not shown). Thus, loss of *Nf1* in post-mitotic PV+ CINs does not recapitulate the phenotypes observed by deletion in developing CINs or progenitors, suggesting that previous phenotypes occur during development/maturation.

**Figure 3:**
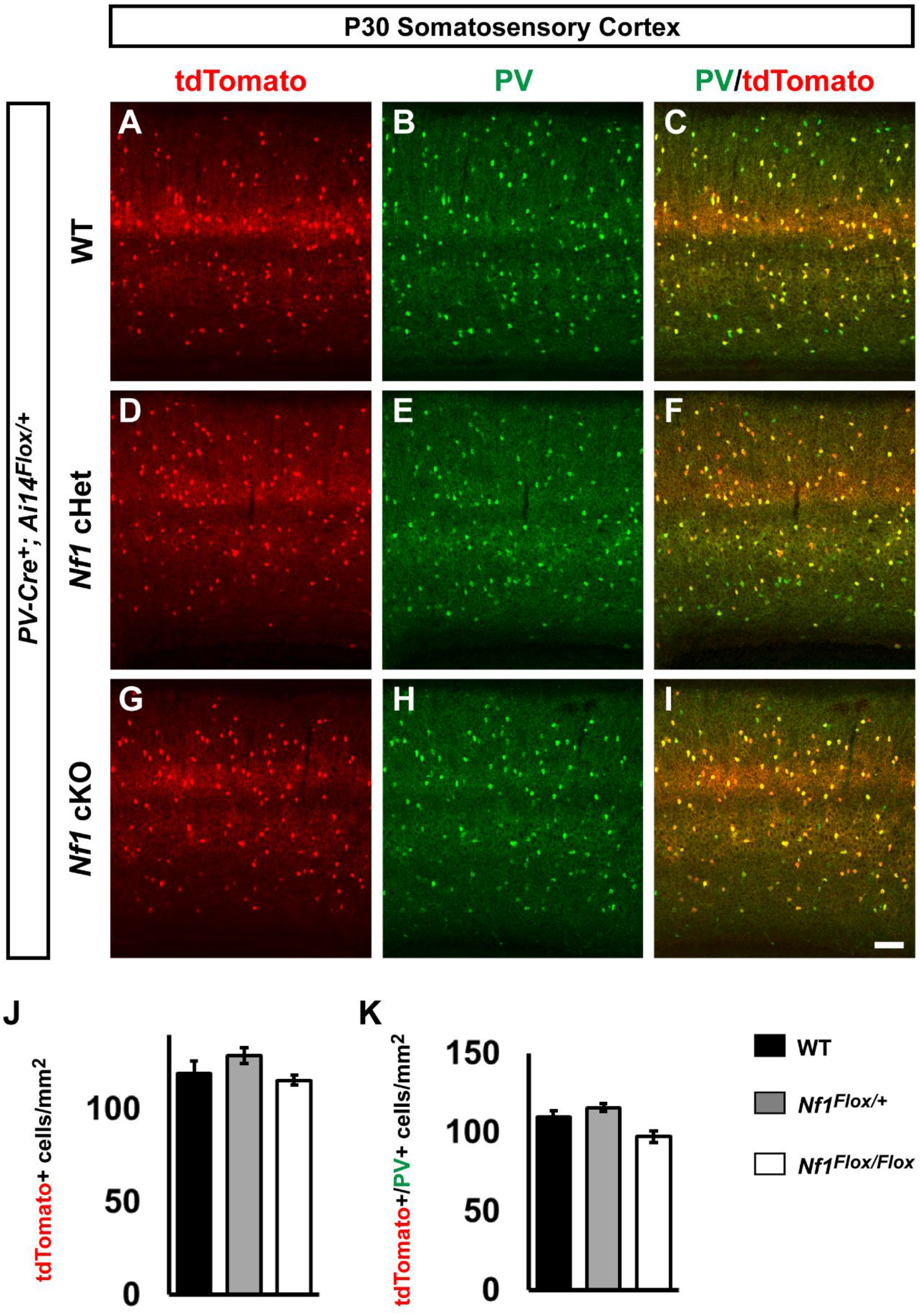
Lack of overt phenotypes after conditional loss of *Nf1* in post-mitotic *PV-Cre* lineages. P30 WT, *Nf1* cHet and *Nf1* cKO coronal issue was visualized for tdTomato, i.e. *PV-Cre*-lineages, (**A, D, G**), PV (**B, E, H**) or merged (**C, F, I**). (**J**) quantification of the cell density of *PV-Cre*-lineages at P30 in the somatosensory cortex. (**K**) quantification of the cell density of tdTomato+/PV+ co-labeled cells. Data are expressed as the mean ± SEM. WT, n = 3, cHet, n = 3 and cKO, n = 4. Scale bar in (**I**) = 100 μm.

### *Nf1* loss in the MGE does not alter cell proliferation

To probe how the loss of PV+ CINs could occur, we assessed whether loss of *Nf1* altered cell proliferation in the MGE. MGE-derived CINs are born throughout mid-gestation, with peak production occurring ~embryonic (E) day 13.5 (Inan et al., 2012; Miyoshi et al., 2007). Moreover, since some PV+ CINs are born later than SST CINs, we also wanted to assess proliferation at E15.5. Thus, we first pulsed pregnant moms with EdU that had either E13.5 or E15.5 embryos for 30 minutes to label proliferating cells in (S) synthesis-phase of the cell cycle, as previous described (Vogt et al., 2015a). We also co-stained these brains for the (M) mitosis-phase marker, phospho-histone 3 (PH3). Labeled cells were counted in both the ventricular zone and subventricular zones of the MGE. This is important, as each of these zones has proliferating cells and whether SST+ vs. PV+ CINs are produced is postulated to be modulated by when cells in these zones become post-mitotic (Petros et al., 2015). However, we did not detect any differences between genotypes for either EdU-pulsed or PH3+ cells at each age (Figure S5A-S5V). Thus, loss of *Nf1* does not alter cell proliferation at these key times of CIN proliferation.

### Loss of PV+ CINs is a cell autonomous consequence of *Nf1* loss

We next asked whether the loss of PV+ CINs was due to *Nf1* loss in CINs or potentially to a non-cell autonomous effect from the immature oligodendrocytes that populate the *Nf1* cKO cortex. Unfortunately, there are no Cre-driver lines to specifically delete *Nf1* in PV-lineage CINs during development. To overcome this, we utilized an MGE transplant assay. This assay takes advantage of the unique migratory properties of CINs, which broadly disperse in the cortex once xenografted (Alvarez-Dolado et al., 2006), while non CINs do not migrate and are only present at the site of injection. This unique feature allowed us to transplant *Nf1* cKO MGE cells, with immature oligodendrocytes remaining in one place while the CINs integrated into the WT cortex of the host mouse, away from the bolus of immature oligodendrocytes (Figures 4A and 4B).

**Figure 4:**
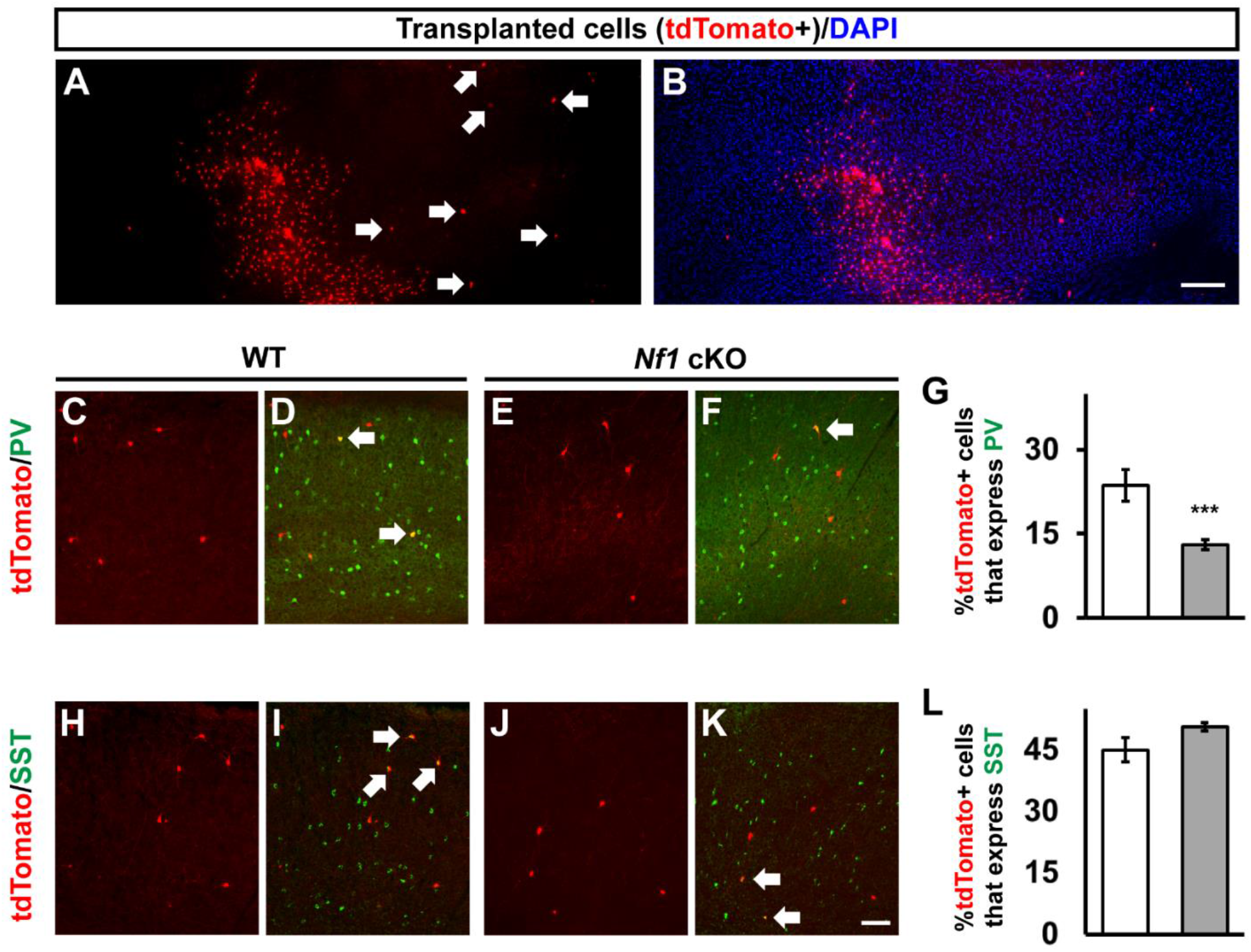
Transplantation of MGE-lineages reveals that the reduction in PV-expressing CINs is cell autonomous. (**A-B**) Transplanted MGE cells (tdTomato+) from *Nf1* cKOs, showing the distinct separation of the small cell body oligodendrocytes and the CINs that migrate away (arrows). (**C-F**) Transplanted MGE cells from either WTs or *Nf1* cKOs (tdTomato+) were co-labeled for PV. (**G**) Quantification of the proportion of tdTomato+ cells that express PV. (**H-K**) Transplanted MGE cells from either WTs or *Nf1* cKOs (tdTomato+) were co-labeled for SST. (**L**) Quantification of the proportion of tdTomato+ cells that express SST. Arrows represent co-labeled cells. Data are expressed as the mean ± SEM. *** p < 0.001. n = 3 transplantations for each group. Scale bars in (**B**) = 200 μm and (**K**) = 100 μm.

To assess whether PV+ CINs from *Nkx2.1-Cre; Ai14; Nf1* cKOs established normal numbers after development, we determined the proportion of transplanted tdTomato+ cells outside of the injection site for PV compared to *Nf1* WT transplanted cells (Figures 4C-4F). A reduction in tdTomato+ cells that expressed PV compared to WT transplanted cells was observed (Figure 4G, p = 0.0007), suggesting that this is a cell autonomous phenotype. Moreover, there was no significant difference in the tdTomato cells that migrated out of the injection site that expressed SST (Figures 4H-4L), consistent with our earlier findings (Figure S2). Overall, these data show that the loss of PV expression is independent of the persistence of MGE-derived oligodendrocytes.

We also wanted to test whether introduction of immature oligodendrocytes would alter the endogenous oligodendrocytes when introduced later in development, i.e. once the *Emx1*-lineage had initially populated the cortex. Thus, we analyzed whether transplanted Olig2+ MGE-lineage cells altered the numbers of later born oligodendrocytes in the cortex from the same transplant experiments as before. When we introduced *Nf1* cKO MGE cells into neonatal cortices, the Olig2+ MGE cells remained in a dense domain at the site of injection (Figures S6A-S6C). Next, we counted the non-tdTomato+ Olig2+ cells in either the injection site or an adjacent region and found that the presence of MGE-lineage Olig2+ cells competed with later born oligodendrocytes in the neocortex, resulting in lower Olig2+ only cells in the transplant region (Figures S6C’, S6C’’ and S6D). This datum suggests that the immature state of the Olig2+ MGE-lineage cells compete with later born cells for positions in the neocortex.

### The *Lhx6* transcription factor is reduced in both *Nf1* cHets and cKOs

Since the *Lhx6* transcription factor is necessary to generate almost all SST+ and PV+ CINs (Liodis et al., 2007; Zhao et al., 2008) and machine learning paradigms predict *Lhx6* to be the primary determinant of MGE-lineage CIN fate (Silberberg et al., 2016), we wanted to assess *Lhx6* expression at P30 in the somatosensory cortex. To this end, we first crossed a *Lhx6-GFP* expressing mouse with our *Nkx2.1-Cre; Nf1^Floxed^* mice to examine MGE-derived CINs in the somatosensory cortex at the same age. We previously had observed that this *Lhx6-GFP* mouse expresses GFP to a greater extent in PV+ MGE-lineage CINs than in SST+ (Pai et al., 2019).

We first assessed the cell density of *Lhx6-GFP*+ CINs in the somatosensory cortex at P30. Interestingly, the number of GFP+ cells were decreased in both the cHets, ~37% reduced vs. WT, and the cKOs, ~77% reduced vs. WT (Figures 5A-5D, WT vs. cHet, p = 0.02 and WT vs. cKO, p = 0.0008, cHet vs. cKO, p = 0.01). Considering there were no changes in SST+ CINs, or any other marker in the cHets, we wanted to assess *Lhx6* expression in another manner. Thus, we performed *in situ* hybridization (ISH) at P30 and counted the number of cells expressing *Lhx6* transcript in the neocortex. Notably, the number of *Lhx6*+ cells was also reduced by half in both the cHets and cKOs (Figures 5E-5H, WT vs. cHet, p = 0.0004 and WT vs. cKO, p = 0.006). While the decrease in total *Lhx6* expression was not as severe as the loss of GFP expression, there was a noticeable reduction in the intensity of *Lhx6* expression in both the cHets and cKOs in CINs that was more severe in the cKOs. Together, these data reveal that loss of *Nf1* results in decreased *Lhx6* levels in a dose-dependent manner.

**Figure 5:**
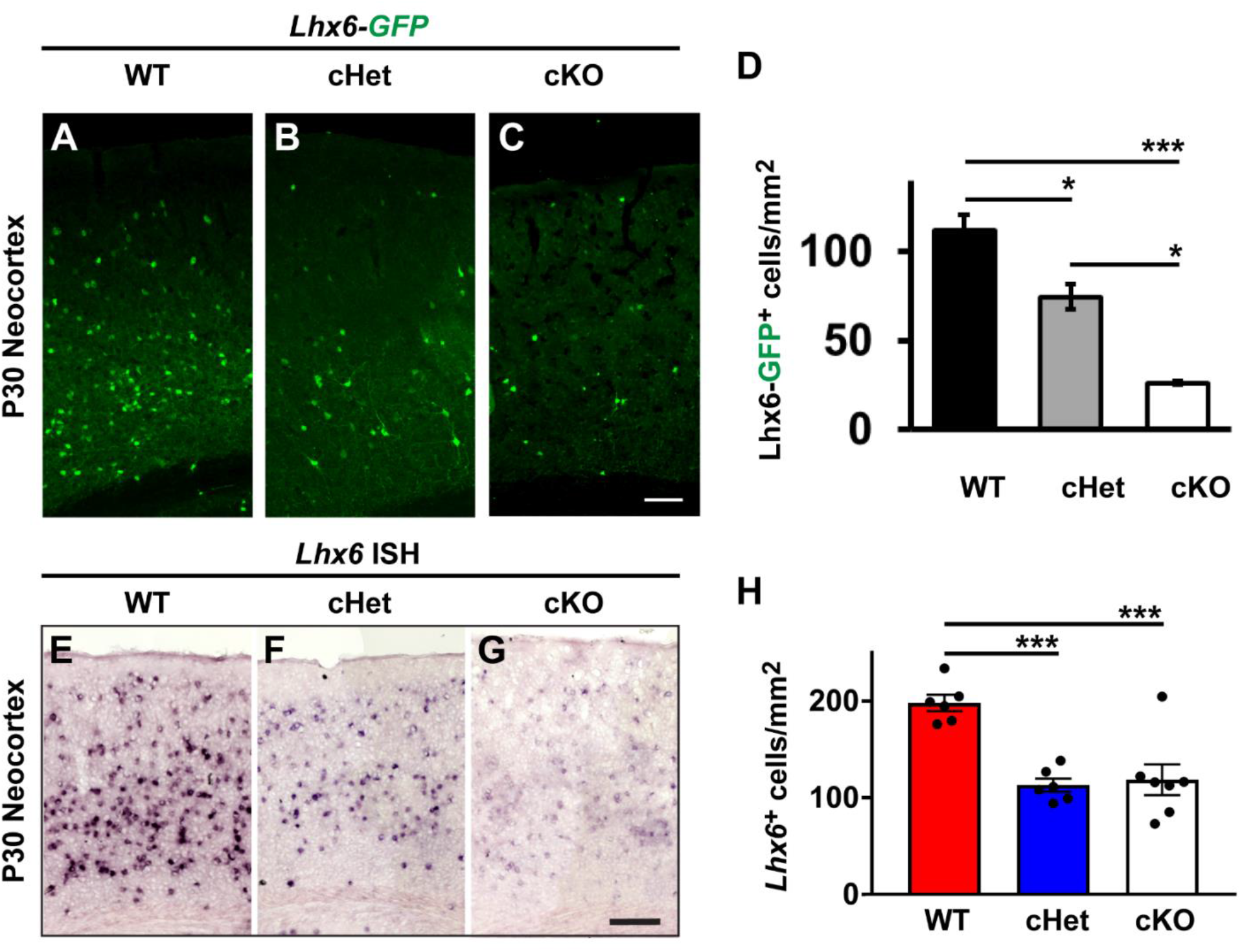
Graded loss of *Lhx6* in *Nf1* cHets and cKOs. P30 WT, cHet and cKO neocortices expressing the *Lhx6-GFP* allele were labelled for GFP (**A-C**). (**D**) Quantification of the *Lhx6-GFP* cell density from each genotype in the neocortex. P30 neocortices were also probed via *in situ* hybridization (ISH) to detect the *Lhx6* transcript (**E-G**). (**H**) Quantification of the number of *Lhx6*+ cells via ISH for each genotype. Data are presented as the mean ± SEM. For Lhx6-GFP counts, n = 3 (WT), n = 6 (cHet) and n = 3 (cKO). * p < 0.05, *** p < 0.001. Scale bars in (**C**) = 100 μm and (**G**) = 200 μm.

### Distinct patterns of *Lhx6* loss during development in *Nf1* cHets and cKOs

To determine if *Lhx6* expression could be responsible for the loss of PV+ CINs, we next examined whether *Lhx6* levels were altered during development. To this end, we first examined *Lhx6-GFP*+ embryos at E15.5, a timepoint when CINs are tangentially migrating through the developing neocortex but just before OLIG2+ cells enter the neocortex (Tekki-Kessaris et al., 2001). The number of Lhx6-GFP+ cells in E15.5 brains had a *Nf1*-loss/dose-dependent decrease (Figures 6A-6C).

**Figure 6:**
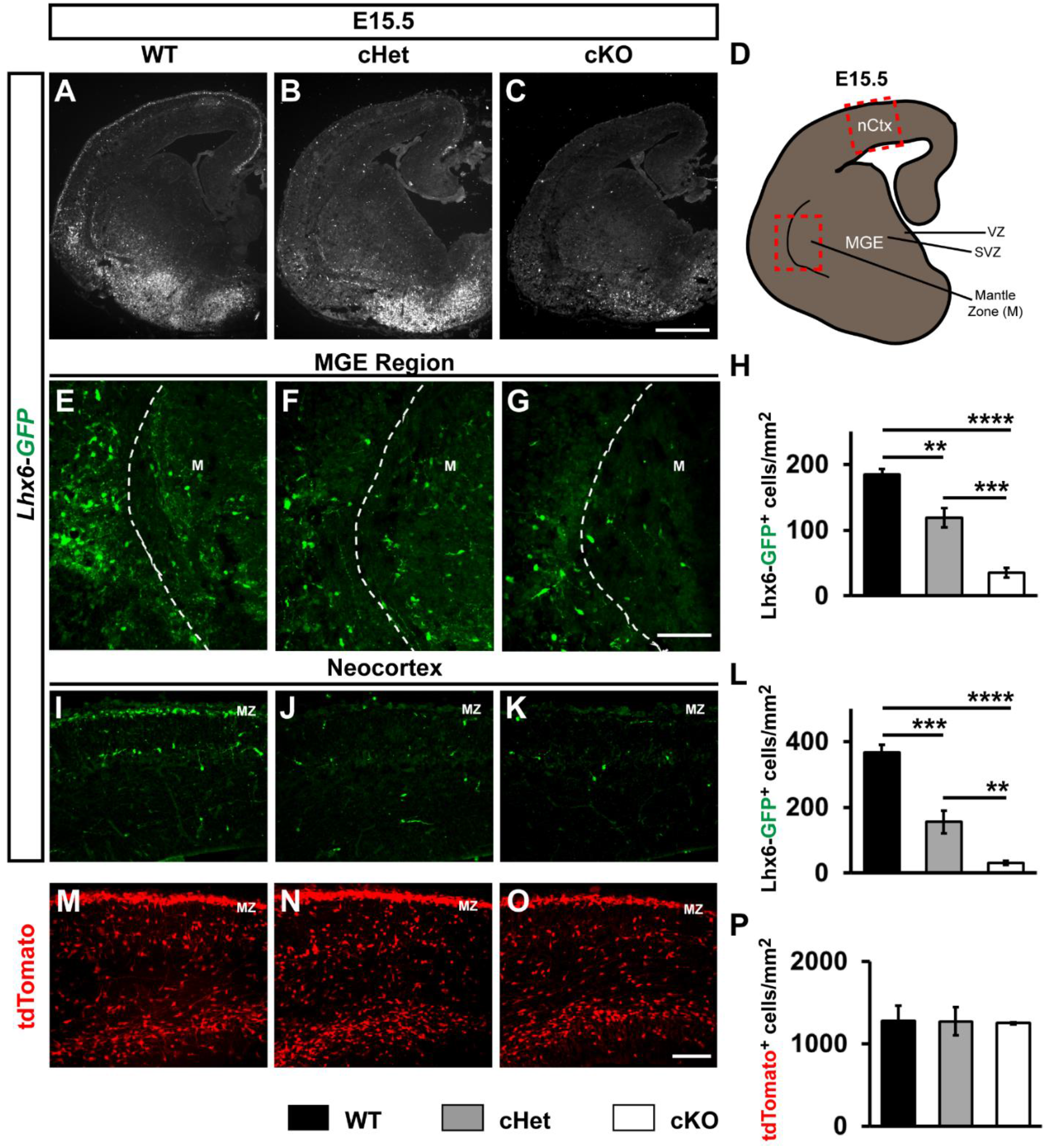
*Lhx6-GFP* cells are depleted in cHets and cKOs by E15.5. E15.5 coronal brain sections were probed for GFP expression from *Lhx6-GFP* expressing mice in WTs, cHets and cKOs (**A-C**). (**D**) Schema of an E15.5 coronal section showing the neocortex region (nCtx) and the regions of the MGE that cells were assessed from. (VZ) ventricular zone and (SVZ) sub ventricular zone. (**E-G**) Lhx6-GFP-labeled images from the mantle zone (M) of the MGE. (**H**) Quantification of the cell density of Lhx6-GFP+ cells in the mantle zone of the MGE. (**I-K**) Tangentially migrating CINs labeled for Lhx6-GFP from the neocortex; marginal zone (mz). (**L**) Quantification of the cell density of Lhx6-GFP+ cells migrating through the neocortex of the MGE. (**M-O**) Tangentially migrating *Nkx2.1-Cre*-lineage cells (tdTomato+) from the neocortex. (**P**) Quantification of the cell density of tdTomato+ cells in the neocortex of the MGE. Data are expressed as the mean ± SEM. n = 3, all groups. ** p < 0.01, *** p < 0.001, **** p < 0.0001. Scale bars in (**C**) = 500 μm, (**G**) and (**O**) = 100 μm.

We asked whether the loss of Lhx6-GFP occurred equally in the MGE mantel zone, where Lhx6-GFP expression is first observed, or if it was more severe in the neocortex, which harbors CINs that have been migrating for several days. To examine this, we first counted Lhx6-GFP+ cells in the mantle zone of the MGE and found that there was already a reduction of these cells in both the cHets, ~40% reduced, and cKOs, ~82% reduced, compared to WTs (Figures 6E-6H, WT vs cHet, p = 0.002, WT vs cKO, p < 0.0001, cHet vs cKO, p = 0.0004). Next, we counted Lhx6-GFP+ cells in the tangentially migrating streams of the neocortex at the same age; these migrating CINs represent populations that are more mature. While not drastic, we did however, notice a continued loss of GFP+ cells in these migrating cells; ~48% reduction in cHets and ~91% reduction in cKOs, vs. WTs (Figures 6I-6L, WT vs cHet, p = 0.0003, WT vs cKO, p < 0.0001, cHet vs cKO, p = 0.009).

While there were reductions in GFP+ cells, there were no decreases in tdTomato+ cells at E15.5 (Figures 6M-6P), suggesting that the loss of *Lhx6* expression was not due to loss of *Nkx2.1-Cre*-lineage CINs. We also examined an early postnatal timepoint, P2, to assess *Lhx6-GFP*+ numbers; there was a consistent decrease in the number of GFP+ CINs in both the cHets and cKOs (Figures S7A-S7D, p < 0.0001 for both groups). Like our data at E15.5, there were no changes in tdTomato+ cells at P2 (Figures S7E-S7H), suggesting that decreased *Lhx6* expression but not cell loss was also occurring at early postnatal ages. Overall, these data show that normal numbers of *Nkx2.1-Cre* lineages migrate into the neocortex during mid-gestation and into early postnatal stages but lose *Lhx6* expression in a *Nf1*-dose and progressive manner.

### Model of *Nf1* dysfunction in MGE-derived oligodendrocytes and CINs

We did not detect a loss *Nkx2.1-Cre*-lineage cell numbers (tdTomato+) embryonically, or in young adult cHets at P30, where accurate counts of CINs could be made, suggesting that in our model loss of *Nf1* may not alter the total number of CINs. However, there is a loss of *Lhx6* that is observed by E15.5 in both the *Nf1* cHets and cKOs and is more severe in the cKOs (Figure 7, left panel). This persists into young adult ages, where we also observe a selective decrease in PV expression in the cKOs (Figure 7, right panel). Notably, while the cHets still have severe reductions of *Lhx6* at P30, they do not have alterations in PV expression; it is possible that enough *Lhx6* is expressed to promote the PV+ CIN cell fate program in cHets but not cKOs. Finally, complete loss of *Nf1* resulted in a persistence of immature oligodendrocytes in young adult brains (Figure 7, right panel). Since *Nkx2.1-Cre* lineages give rise to both MGE-derived oligodendrocytes and CINs, it may be possible that oligodendrocyte progenitors outcompete CIN progenitors in the MGE, as was observed in another *Nf1* model (Wang et al., 2012a). However, using a novel transplantation approach, where oligodendrocytes can be physically separated from CINs, we found that each phenotype was cell autonomous (Figure 4). Moreover, we did not detect any changes in proliferation in the progenitor domains of the MGE that would suggest otherwise (Figure S5). Thus, we propose that oligodendrocyte and CIN phenotypes are independent indices of *Nf1* dysfunction.

**Figure 7:**
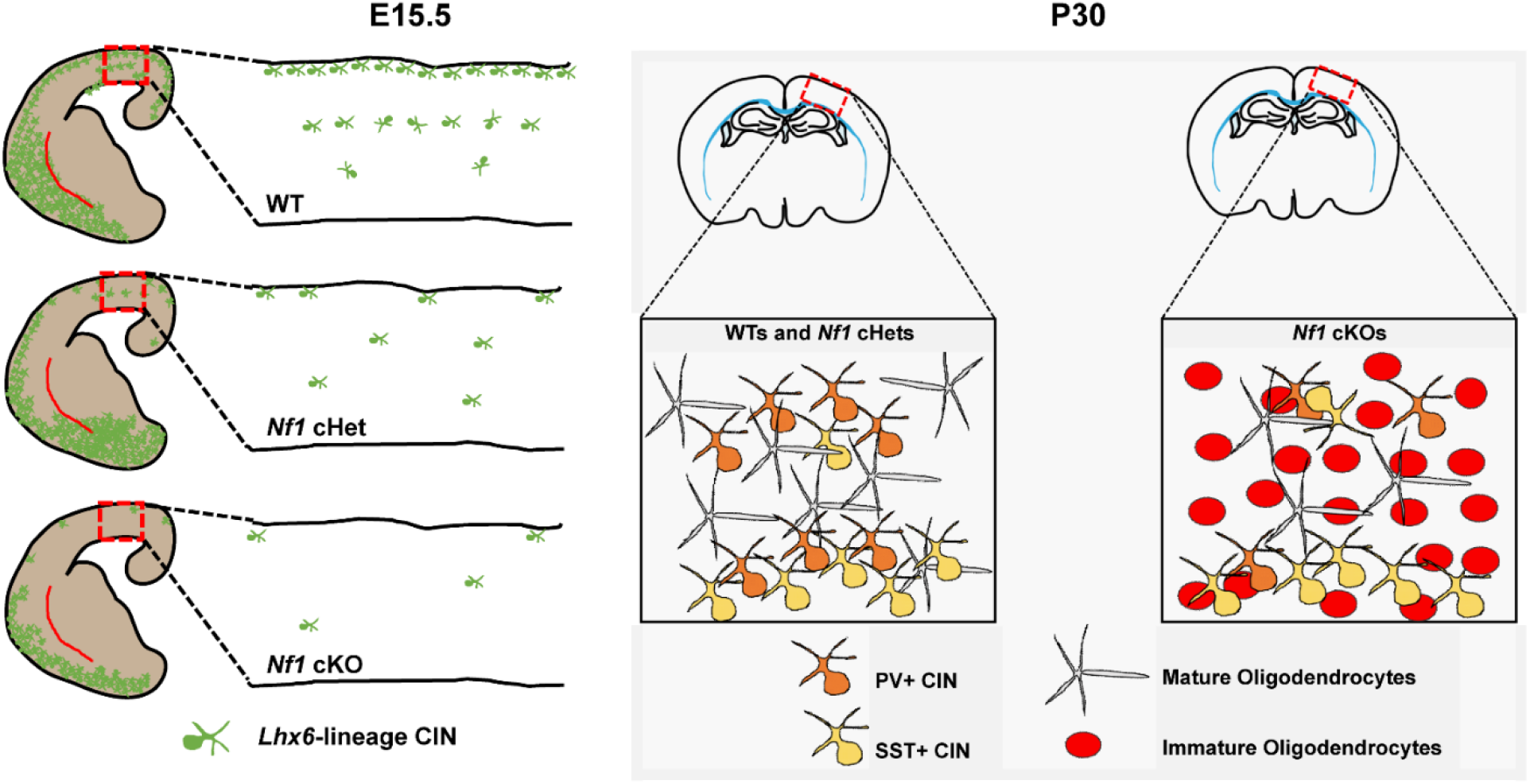
Phenotypic model of *Nf1* loss in CIN progenitors. (**Left panel**) After conditional loss of *Nf1* in *Nkx2.1-Cre*-lineages, *Lhx6* expression is decreased in a dose-dependent fashion in *Nf1* cHets and cKOs. Boxes and insets show the expression of *Lhx6* as these cells develop and migrate through the neocortex. (**Right panel**) By young adult ages, *Nf1* cHets still exhibit a loss of *Lhx6* (not shown herein) but generate normal numbers of SST+ and PV+ CINs, suggesting that sufficient amounts of this critical transcription factor were present to instruct their development. However, in *Nf1* cKOs, which have drastic reductions in *Lhx6* expression, there is a specific reduction in PV expression without any change in the SST+ CIN group. This suggests that either a critical dosage of *Lhx6* or a developmental window are insufficient/lost during the development of PV+ CINs in *Nf1* cKOs. Finally, *Nf1* cKOs also have a persistence of immature oligodendrocytes derived from *Nkx2.1-Cre*-lineages. We determined that both this phenotype and the loss of PV expression are independent events, using a novel cell transplantation approach. Notably, these immature oligodendrocytes outcompete later born oligodendrocytes for occupancy in the neocortex and may also contribute to symptoms in NF-1.

## Discussion

The increased prevalence of ASD and learning disabilities in many of those diagnosed with NF-1 (Garg et al., 2013; Morris et al., 2016) likely have a cellular and molecular basis. Herein, we explored how cells derived from an embryonic progenitor domain, the MGE, which generates both oligodendrocytes and GABAergic CINs, are altered after loss of *Nf1* in mice. This was an important question, as both GABAergic CINs and oligodendrocytes have been implicated in NF-1 (Bennett et al., 2003; Costa et al., 2002; Cui et al., 2008; Lee et al., 2010; Mayes et al., 2013; Wang et al., 2012a), yet how these populations from the MGE are affected had not been explored.

We first discovered a glial phenotype that was different than the competition between gliogenesis and neurogenesis previously reported in a different region of the brain (SVZ of the dorsal LGE), where progenitors preferentially generated oligodendrocytes at the expense of neurons (Wang et al., 2012a). In our study, MGE-derived oligodendrocytes that lacked *Nf1* expressed the immature marker, NG2, and persisted into adult stages, thereby competing out later-born oligodendrocytes. However, there were no alterations in the number of MGE progenitors generated and normal number of CINs were found in the embryonic neocortex. This is interesting, as ventral-derived oligodendrocytes of the spinal cord outnumber their later-born dorsal counterparts and loss of *Nf1* in the spinal cord results in elevated immature markers (Bennett et al., 2003; Lee et al., 2010), similar to what we observe in MGE-derived oligodendrocytes that also have ventral origins. Moreover, it suggests that early born oligodendrocytes can non-cell autonomously influence the development of their later born counterparts, suggesting a form of glial communication during brain development that may help us understand how developing oligodendrocytes establish proper numbers in the adult nervous system.

The excitation/inhibition (E/I) hypothesis was originally put forward (Rubenstein and Merzenich, 2003) as one postulate for how some symptoms of ASD arise, and was recently updated (Sohal and Rubenstein, 2019). Alterations in the neocortical excitation/inhibition balance (Canitano and Pallagrosi, 2017; Gao and Penzes, 2015; Uzunova et al., 2016; Yizhar et al., 2011) and GABAergic signaling, have been implicated in ASD (Costa et al., 2002; Hussman, 2001; Robertson et al., 2016). Previously, increased GABA release was proposed to modulate hippocampal LTP and learning following deletion of *Nf1* using a pan-GABAergic Cre line (Cui et al., 2008). Our study demonstrates that loss of *Nf1* in MGE-derived CINs, a deletion more spatially restrictive than the previous report, results in a specific loss of PV expression. This is interesting, as the numbers of PV+ are uniquely reduced in the prefrontal cortices of humans with ASD, compared to other CIN types (Hashemi et al., 2017). Thus, manipulating PV+ CIN function and numbers may be potential therapeutic inroads in the future.

While our deletion strategy led to a loss of *Lhx6* and PV expression, future studies are needed to determine what the net effect on excitatory/inhibitory balance is. However, some clues have come from another research group that found a loss of inhibition using similar techniques (Holter et al., co-submitted manuscript), which is at odds with previous reports on *Nf1* loss using a broader deletion strategy (Cui et al., 2008). This raises an interesting point, which is that while our more restrictive manipulation may reveal an important role for PV+ CIN development, there may be other GABAergic neurons responsible for the increased GABAergic tone previously reported (Cui et al., 2008). Notably, we found that *Nf1* transcripts were higher in CGE-lineage CINs, a group that our studies did not manipulate herein, but were included in the Cui et al studies. Assessment of these cells could be a valuable inroad to further understanding how NF-1 cognitive symptoms arise.

Both *Nkx2.1* and *Lhx6* regulate distinct conserved DNA regulatory elements to determine MGE regional identity and initiate the expression of specific genes in developing CINs (Sandberg et al., 2016). *Lhx6* expression in MGE-derived CINs is indispensable for their normal tangential and radial migration, survival and specification of molecular identity (Elbert et al., 2019; Liodis et al., 2007; Neves et al., 2013; Vogt et al., 2014; Zhao et al., 2008). Finally, the expression of *Lhx6* is repressed in several types of cancers via hypermethylation of its regulatory elements (Estécio et al., 2006; Jung et al., 2010, 2011; Liu et al., 2013). While the signaling events repressing *Lhx6* are not known, it is possible that the RAS oncogene mediates these effects and could potentially lead to *Lhx6* repression in CINs. In the light of these evidences, we were interested in assessing how this important control gene was impacted by loss of *Nf1*.

While there were no differences in the numbers of *Nkx2-1-Cre* lineage cells that migrated into the neocortex at embryonic ages, we observed a progressive loss of *Lhx6* in a *Nf1*-dose dependent manner. Contrary to earlier studies (Liodis et al., 2007; Neves et al., 2013; Zhao et al., 2008), the selective loss of PV+ CINs and the preservation of the SST+ CIN numbers in our mutants raises important questions regarding the role of *Lhx6* in the regulation of CIN phenotypes and the pathology of NF-1. One possibility is that the loss of the NF1 protein was delayed following depletion of the gene, thus allowing for MGE cells that were born early to maintain *Nf1* and *Lhx6* expression in some portions of the MGE. Notably, *Lhx6* hypomorphs had a selective decrease in the emergence of SST+ CINs and no changes in PV+ CINs (Neves et al., 2013); of note, these mice had reduced *Lhx6* expression at every stage of development, while our conditional mutants needed time to delete the *Nf1* gene, wait for the protein to degrade and then induce the effect on *Lhx6*. Taken together, this sets up an interesting hypothesis into how MGE-lineage CIN fate may be attained. Potentially, *Lhx6* is necessary at very early stages to establish the generation of the SST-lineage but may need to be expressed at some critical threshold, i.e. below what Neves et al., 2013 showed, i.e. ~60% reduction, but above the 90% reduction in the *Nf1* cKOs observed herein, for PV-lineages to be established. This may be inline with a recent report proposing that the SST-lineage may preferentially be determined very early in the VZ of the MGE while PV-lineages are more likely to be programmed later, after becoming postmitotic in the SVZ of the MGE (Petros et al., 2015) and that chromatin may be sustained in a manner for PV expression to occur during adolescence, given some critical signal. Since our model demonstrates a progressive loss of *Lhx6* in *Nf1* mutants, it sets up a unique system in which the very early stages are preserved and the progressive developmental stages lose *Lhx6*, thus setting up a situation where later stages of development can be assayed for the effects of *Lhx6* loss. This situation leads to a drastic loss of PV expression but no change in SST expression.

Our data reveal new insights into how *Nf1* regulates the development of MGE-derived oligodendrocytes and CINs. However, *Nf1* controls RAS/MAPK signaling from an initial stage and whether other RASopathy genes may influence oligodendrocyte and CIN development is an ongoing question. Moreover, *Nf1* can also regulate adenylyl cyclase and MTOR signaling pathways (Johannessen et al., 2005; Tong et al., 2002). Interestingly, another group found that a constitutively active MEK1 mouse model also exhibits a similar loss of PV+ CINs in the neocortex (Holter et al., co-submitted manuscript). This observation suggests that PV+ CIN development/maturation may be regulated through the core RAS/MAPK pathway and may not be due to other *Nf1*-regulated signaling events. It is also interesting the Holter et al. see an embryonic increase in CIN apoptosis while we don’t observe changes by E15.5. However, we think these differences may arise from our different mouse models, i.e. we utilized a loss of function while they used a gain of function approach. The loss of *Nf1* function may require more time to exert a phenotype, i.e. the protein and mRNA need to be degraded after Cre-mediated deletion while the gain of function approach they used is on in MGE progenitors.

The persistence of immature oligodendrocytes and loss of PV+ CINs in *Nf1* mutants opens new avenues for probing the mechanisms underlying RAS/MAPK regulation of these important developmental events. The MEK inhibitors that cross the blood brain barrier such as SL327 (Satoh et al., 2011) have been effectively employed as a single-dose intraperitoneal administration at P6 to induce apoptosis in cortical neurons and oligodendrocytes (Yufune et al., 2015). Interestingly, the abnormal gliogenesis in lieu of neurogenesis during the developmental trajectory in a germline RAS/MAPK overactivation model was therapeutically reversed with transient intervention with a MEK inhibitor during neonatal stages (Wang et al., 2012b). Our studies provide novel insights into the molecular and cellular alterations that may underly some cognitive symptoms in those diagnosed with NF-1. Moreover, these new findings may have broader implication for other RASopathies and associated neurodevelopmental disorders, such as ASD.

### Experimental procedures

#### Animals

All mouse strains have been published: *Ai14* Cre-reporter (Madisen et al., 2010), *Nkx2.1-Cre* (Xu et al., 2008b), *Lhx6-GFP* (Gensat), *Nf1^flox^* (Zhu et al., 2001)*, PV-Cre* (Hippenmeyer et al., 2005). *Nf1^flox^* mice were initially on a mixed C57BL6/J, CD-1 background, then backcrossed to CD-1 for at least four generations before analysis. For timed pregnancies, noon on the day of the vaginal plug was counted as embryonic day 0.5. All animal care and procedures were performed according to the Michigan State University and University of California San Francisco Laboratory Animal Research Center guidelines.

#### Cell counting and statistical analysis

Detailed methods can be found in extended experimental procedures.

#### Immunofluorescent tissue staining

Brains were harvested either from embryos and then fixed overnight in 4% paraformaldehyde (PFA) or from postnatal mice that were perfused with saline followed by 4% PFA; the latter group was post-fixed in 4% PFA for 30 minutes. The brains were then sunk in 30% sucrose, embedded in OCT and sectioned using a cryostat (Tissue-Tek Cryo-3). Immunofluorescent labeling was performed on 25 μm (P2 and older) or 20 μm (embryonic) cryosections with the primary antibodies listed in the Star methods. The appropriate 488, 594 or 647 Alexa-conjugated secondary antibodies were from Thermo Fisher Scientific. Sections were cover slipped with Vectashield containing DAPI (Vector labs).

#### *In situ* hybridization

For standard *in situ* hybridization, a *Lhx6* Digoxigenin-labeled riboprobe was used as previously described (Long et al., 2009). *In situ* RNA hybridization was performed as previously described (Cobos et al., 2007). Imaging was done with 60X objective (Nikon Apo 1.4 oil) under Nikon Ti microscope with DS-Qi2 color camera.

#### MGE transplantation

MGE transplantations were done as previously described (Vogt et al., 2014, 2015b). A detailed description can be found in extended experimental procedures.

## Supporting information

Supplemental data and resources

## Author contributions

**KA** and **ELLP** performed most experiments, conceived ideas and analyzed data. **SMB** performed cell counts. **AMS** performed EdU, PH3 staining and cell measurements. **JTN** performed EdU and PH3 counts. **AP** calculated *Nf1* transcript levels in single CINs. **KA, ELLP** and **DV** conceived ideas and wrote the manuscript. All authors read and contributed to writing the manuscript.

## Acknowledgements

**ELLP** was funded by UCSF neuroscience graduate program and NIMH R01 MH081880. **JLR** was funded by Nina Ireland, NIMH R01 MH081880 and NIMH R37/R01 MH049428. DV was funded by a pilot grant from CTSI (#1111111). **KA**, **AMS**, **JTN** and **DV** were supported by the Spectrum Health-MSU Alliance Corporation.

