## Supplemental data and resources for "*Nf1* deletion results in depletion of the *Lhx6* transcription factor and a specific loss of parvalbumin+ cortical interneurons"

### Contact for Reagent and Resource Sharing

### Experimental Model and Subject Details

Mouse lines

All mice strains have been published: *Ai14* Cre-reporter (Madisen et al., 2010), *Lhx6-GFP* (GENSAT) Tg(Lhx6-EGFP)BP221Gsat, *Nf1*$^{flox}$ (Zhu et al., 2001), *Nkx2.1-Cre* (Xu et al., 2008) and *PV-Cre* (Hippenmeyer et al., 2005). *Nf1*$^{flox}$ mice were initially on a mixed C57BL6/J, CD-1 background, then backcrossed to CD-1 for at least four generations before analysis. For timed pregnancies, noon on the day of the vaginal plug was counted as embryonic day 0.5. Both male and female mice were assessed. We did not observe any gross differences between genders and combined all data for each genotype. All animal care and procedures were performed according to the Michigan State University and University of California San Francisco Laboratory Animal Research Center guidelines.

### Method Details

Cell Counting

We prepared either 20 µm thick cryo-sectioned coronal tissues, for embryonic ages, or 25 µm sections for all postnatal ages. To determine cell density (cells/mm$^2$), we counted the number of cells in a given section from the neocortex, hippocampus or striatum and then divided by the area of that region. To calculate the % of tdTomato$^+$ cells that co-labeled with specific markers, we divided the number of co-labeled cells by the total number of tdTomato$^+$ cells. For cell transplants, all tdTomato$^+$ cells that were assessed for PV or SST were counted in the neocortex from all sections in a rostral to caudal series that were outside of the injection sites where oligodendrocytes resided. For quantification of OLIG2+ cells in transplantation experiments, boxes were drawn in Image-J within the transplant site and an adjacent site that lacked transplanted cells. OLIG2+/tdTomato- cells were then counted in each box and divided by the box area to determine the cell density. For soma size quantification, the perimeter of each tdTomato$^+$ cell's soma was traced in Image J, and at least 25 cells were measured and averaged for each (n) reported.

EdU labeling

Pregnant mice with either E13.5 or E15.5 embryos were pulsed with EdU (10mg/ml) at a dose of 50mg Edu/kg body weight as previously described (Vogt et al., 2015). After 30 minutes, mice were sacrificed, and embryos collected in ice-cold PBS (pH 7.2). Embryos were put in 4% PFA and fixed overnight at 4°C and sunk in 30% sucrose before embedding in OCT. Edu$^+$ cells were visualized by following standard procedures in the Clik-iT EdU plus kit (Thermo Fischer) and then co-stained with antibodies before being labeled with DAPI.

### Fluorescent in situ hybridization (FISH)

FISH was performed according to (Duan et al., 2018). Brain tissue from *Nkx2.1-Cre*-lineage P30 WT and *Nf1* cKOs were perfused and incubated with 4% PFA for 1 hour, following by 30% sucrose/PBS cryoprotection until the day of section. Tissues were embedded in OCT and cryo-sectioned to generate 40um tissue sections. To generate the *Nf1* DNA vector and riboprobe, *Nf1* cDNA was PCR amplified from a homemade mouse cDNA library synthesized from P0 neocortex using Superscript II. The following primers are used: 5' GAG AAT CGA TCC CTC ACA GCT TCG AAG TGT and 3' ATA TTC TAG AGG ACC CAG ATA CGC GAG AAG. ClaI and XbaI restriction enzymes sites were introduced (underlined). The primers target the *Nf1* floxed region (exons 31 and 32). Next, the *Nf1* PCR product and the vector, pSP73 (Promega Cat # P2221), were digested with ClaI and XbaI, and then ligated. The *Nf1* RNA anti-sense fluorescein-labeled probe was generated by T7 RNA polymerase (Roche) and a Fluorescein labeling kit (Roche) from a NdeI linearized vector; size of the probe was 572bp. Imaging was done with 60X objective (Nikon Apo 1.4 oil) under Nikon Ti micsroscope with DS-Qi2 color camera.

### MGE cell transplantation

E13.5 MGEs from individual *Nkx2.1-Cre; Ai14* embryos that were either *Nf1* WT or *Nf1* Flox/Flox were dissected in ice-cold HBSS and then kept on ice in DMEM media (containing 10% fetal bovine serum). MGEs were then mechanically dissociated with a p1000 pipette tip. Cells were then pelleted in a tabletop centrifuge at low speed (700xg, ~4 minutes). The final pellet was left under a few µls of media, put on ice, and the remaining media was removed with a fine point kim wipe before the injection needle was loaded. For injections, a glass micropipette with a 45º beveled tip, ~50 µm outer diameter, was preloaded with sterile mineral oil and cells were then front-loaded into the tip of the needle using a plunger connected to a hydraulic drive (Narishige MO-10). The moveable part of the drive that controlled the plunger was mounted to a manual micromanipulator (M3301R, World Precision Instruments), which aligned the plunger with the glass needle and allowed for movement of the cells in and out of the needle. Pups were anesthetized on ice for 1-2 minutes before being placed on a mold for injections. Each pup received 3-6 injections of cells, at 70 nl per site, in their right hemisphere. These sites were about 1 mm apart from rostral to caudal and medial to lateral and were then injected into layers V-VI of a P1 WT neocortex. After injections, pups recovered and were then put back with their mother. The transplanted mice were sacrificed at 35 days post-transplant and transcardially perfused with PBS followed by 4% PFA. Brains were then post-fixed in 4% PFA for a short interval, ~30 minutes, and then sunk in 30% sucrose before embedding in OCT.

Single CIN *Nf1* RNA transcript assessment

The unique transcripts per million for different CIN groups and subgroups was determined using the previously described dataset (Paul et al., 2017). Briefly, the unique *Nf1* transcripts per million were assessed from a combination of Cre, Flp and the appropriate reporter mice in CINs from young adult mice. The CIN types probed were chandelier, parvalbumin-lineage, somatostatin-lineage that were either nitric oxide synthase-1 or calretinin expressing as well as vasoactive intestinal peptide-lineages that were either calretinin or cholecystokinin expressing.

**Quantification and statistical analysis**

For anatomical analyses, statistics were performed using Prism version 6, a p value of $< 0.05$ was considered significant. For all parametric measurements of three or more groups, we used a One-Way ANOVA with either Bonferroni or Tukey post test to determine significance. For parametric measures of two groups, a two-tailed T-test was performed. For non-parametric data sets, we used a Chi-squared test to determine significance.

**Supplemental Figure 1**

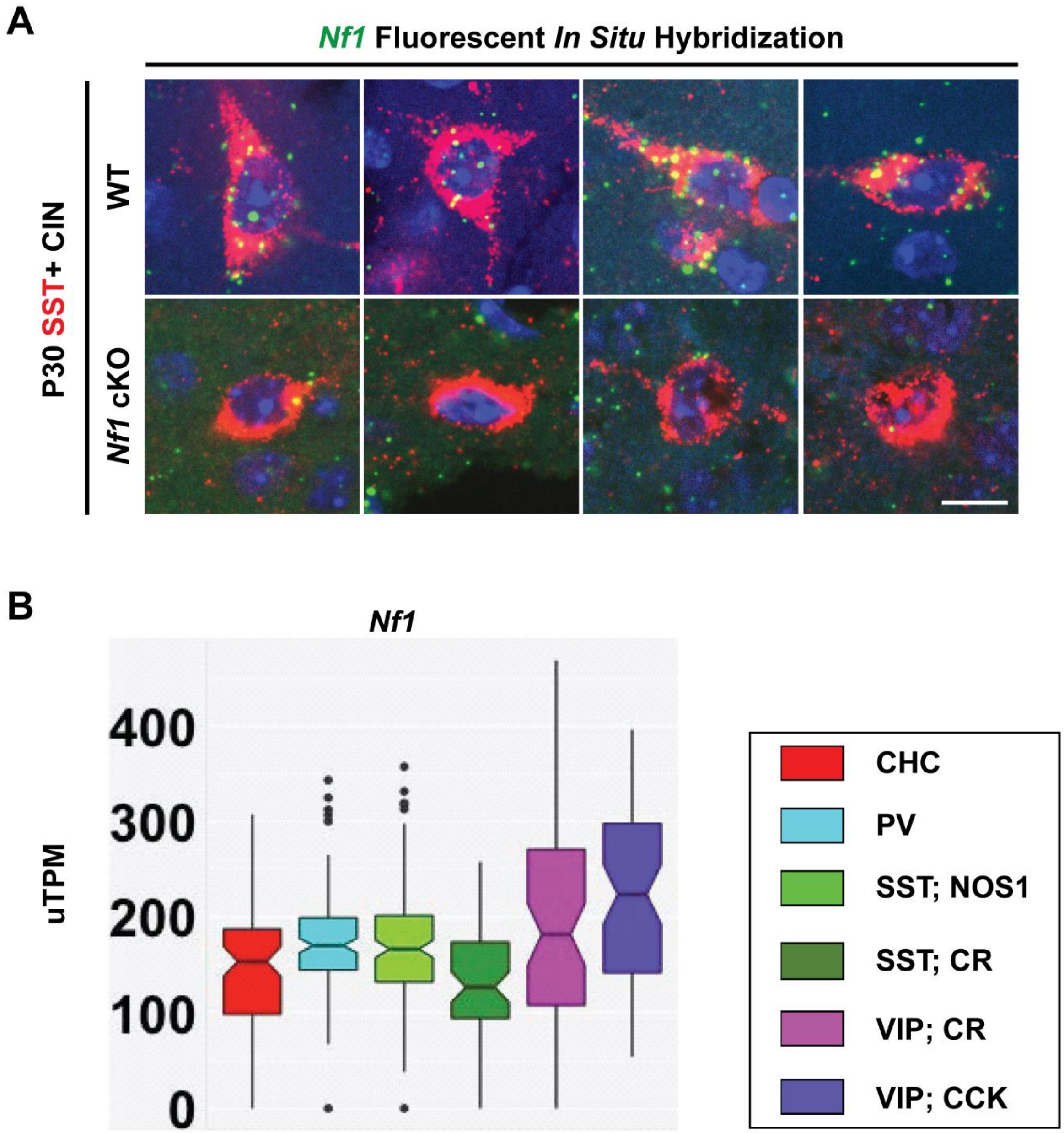


**Supplemental Figure 1: *Nf1* transcript is found in most CIN classes and validation of transcript loss in *Nkx2.1-Cre*-lineage SST+ CINs in adult cortex.**

(**A**) Fluorescent *in situ* hybridization (FISH) for the floxed *Nf1* exon in P30 WT (top panels) and *Nf1* cKOs (bottom panels) in the somatosensory cortex. *Nf1* transcript was co-stained with SST to verify expression and loss in MGE-derived CINs. *Nf1* transcript levels from purified single CINs from P28-P35 cortex using distinct Cre-driver lines (**B**). Box plots show quantification of the unique transcripts per million (uTPM) from single CIN cells. Single dots represent outlier values. (CHC) chandelier cell, (PV) parvalbumin, (SST) somatostatin (NOS1) nitric oxide synthase, (CR) calretinin, (VIP) vasoactive intestinal peptide, (CCK) cholecystokinin. Scale bar in (**A**) = 20 µm.

**Supplemental Figure 2**

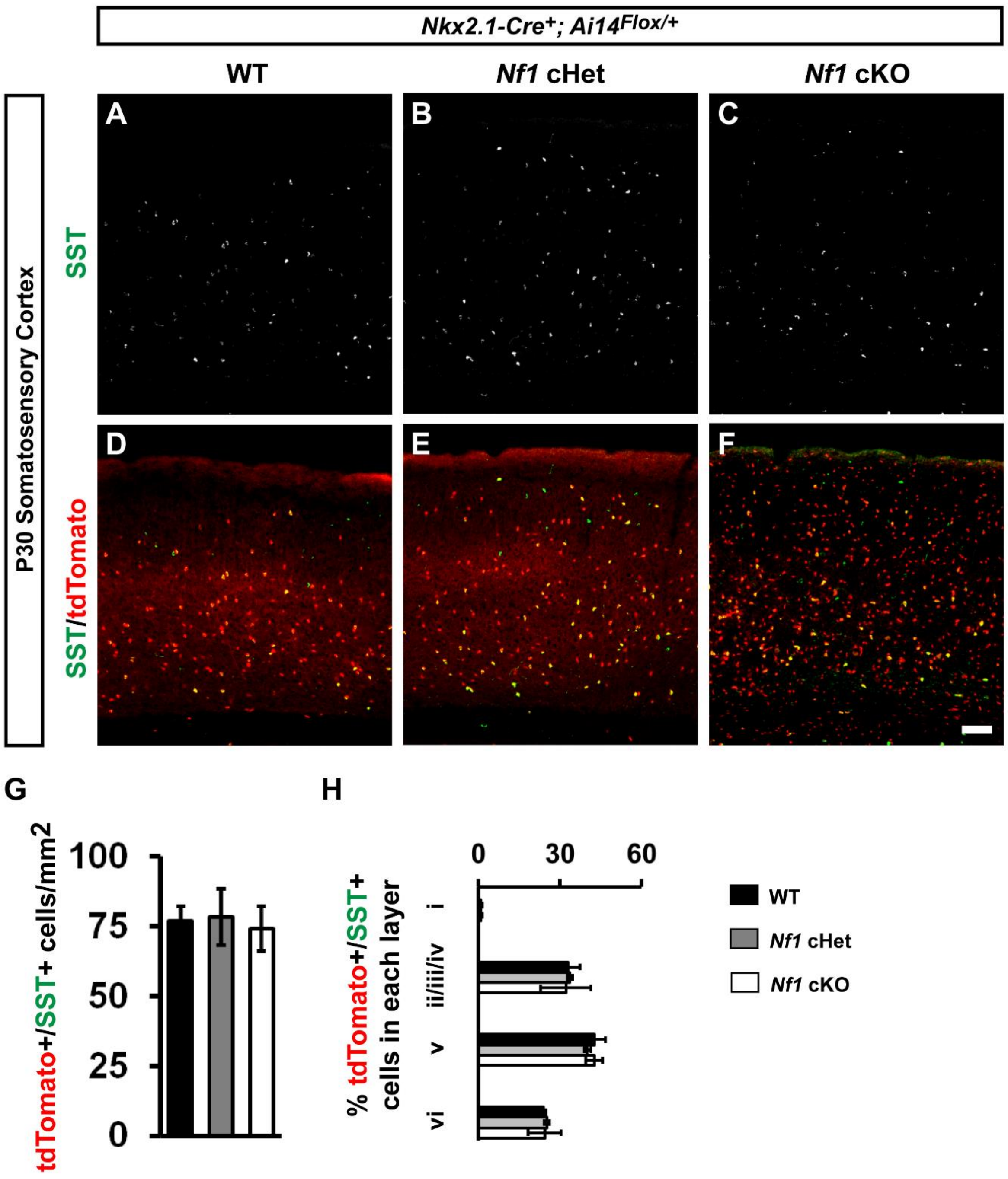


**Supplemental Figure 2: No changes in SST+ CINs after loss of *Nf1*.**

P30 somatosensory cortices from WTs, cHets and cKOs were labeled for SST (**A-C**) and merged with tdTomato+ *Nkx2.1-Cre* lineages (**D-F**). (**G**) Quantification of the cell density of tdTomato+/SST+ CINs in the somatosensory cortex at P30. (**H**) Soma size quantification of tdTomato+/SST+ CINs; (AU) arbitrary units. (**I**) Quantification of the proportion of tdTomato+/SST+ cells in each cortical lamina. Data are expressed as the mean ± SEM. All groups, n = 4. Scale bar in (**F**) = 100 µm.

**Supplemental Figure 3**

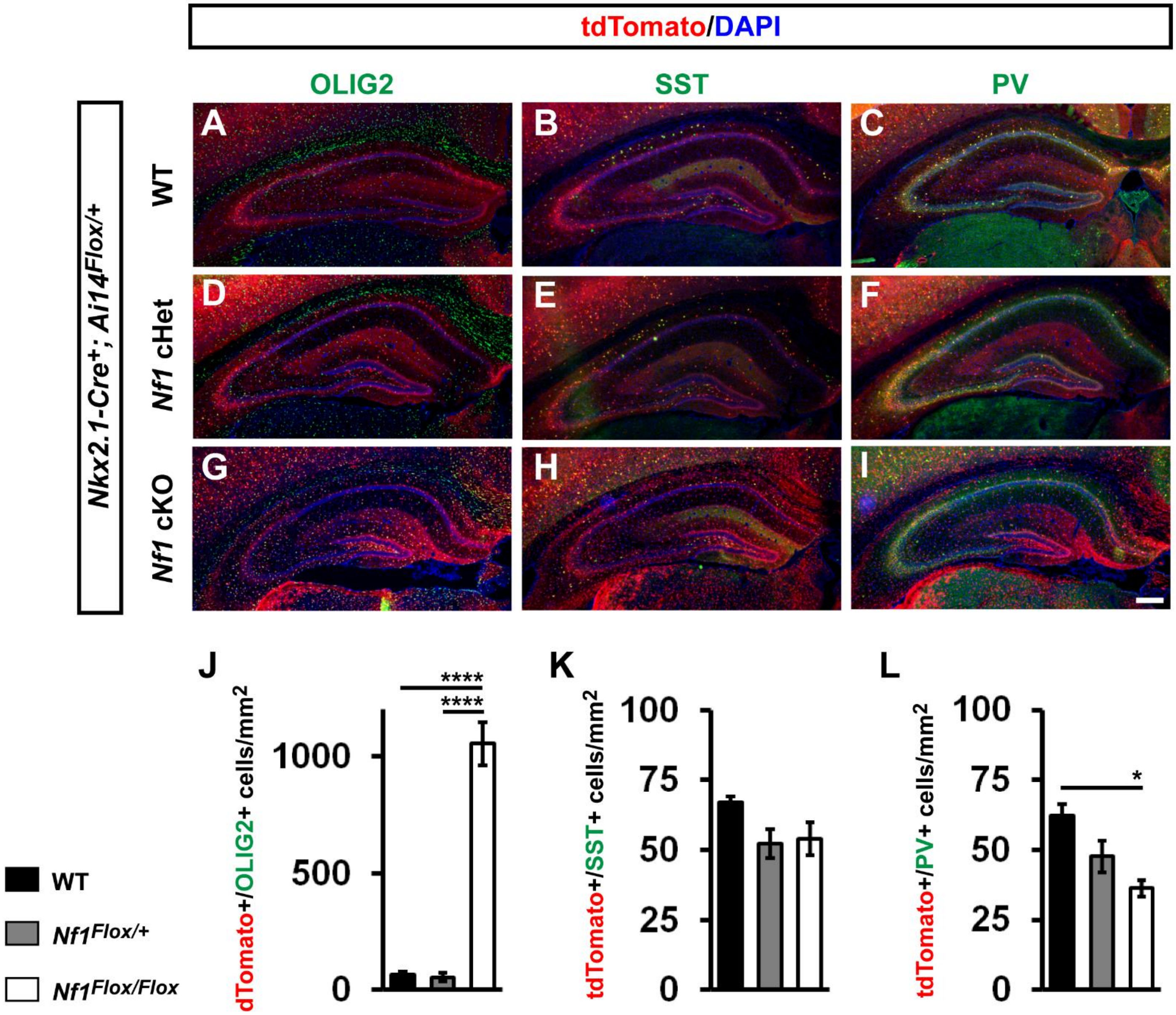


**Supplemental Figure 3: *Nkx2.1-Cre* conditional loss of *Nf1* leads to an increase in immature oligodendrocytes and a loss of PV+ CINs in the hippocampus.**

P30 WT (**A-C**), *Nf1* cHet (**D-F**) and *Nf1* cKO (**G-I**) coronal tissue was visualized for tdTomato (*Nkx2.1-Cre*-lineages) and either Olig2, SST or PV. (**J**) Quantification of co-labeled tdTomato/OLIG2+ cell density. (**K**) Quantification of co-labeled tdTomato/SST+ cell density. (**L**) Quantification of co-labeled tdTomato/PV+ cell density. Data are expressed as the mean ± SEM. All groups, n = 3. * $P < 0.05$, **** $p < 0.0001$. Scale bar in (**I**) = 100 µm.

**Supplemental Figure 4**

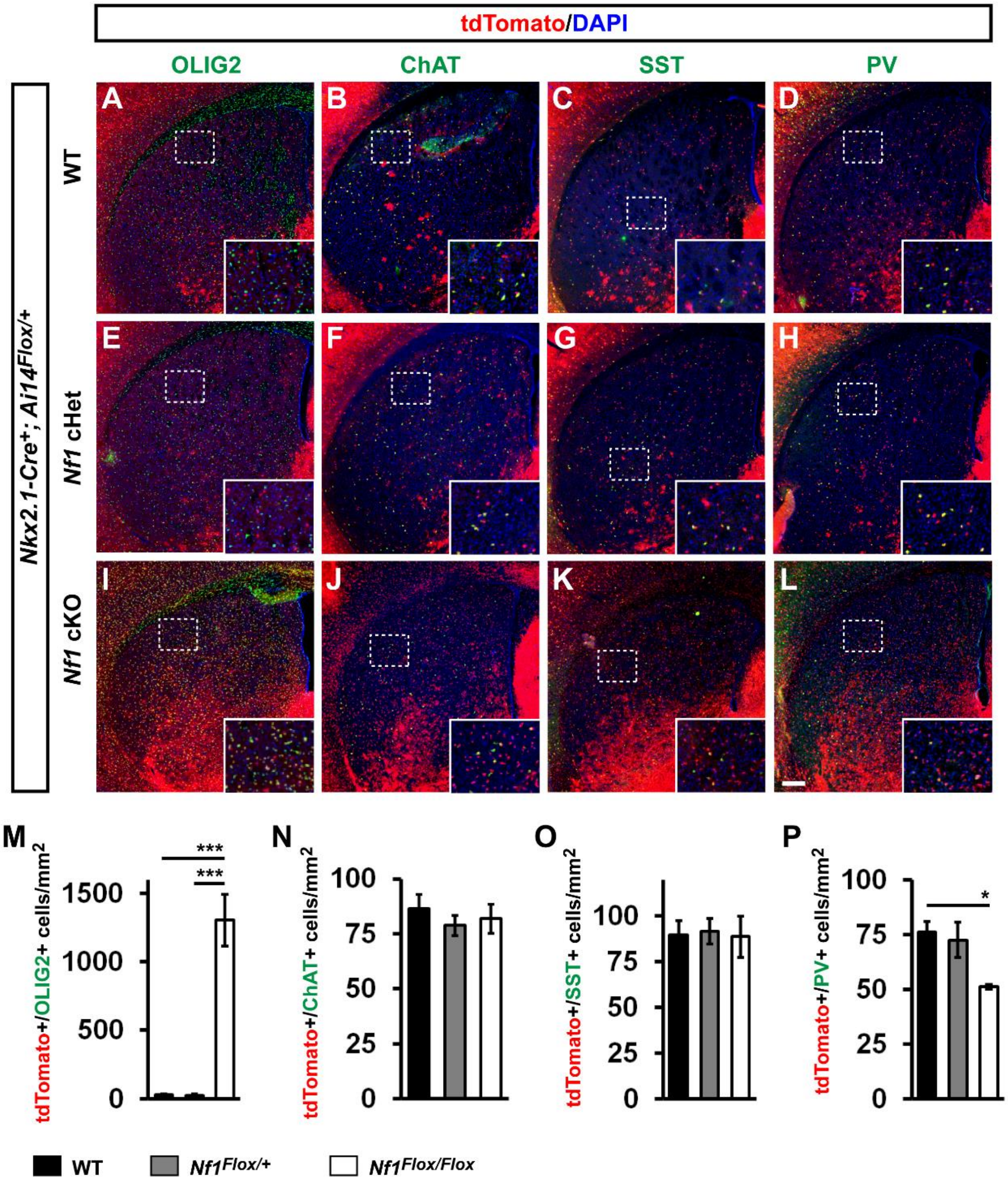


**Supplemental Figure 4: *Nkx2.1-Cre* conditional loss of *Nf1* leads to an increase in immature oligodendrocytes and a loss of PV+ CINs in the striatum.**

P30 WT, *Nf1* cHet and *Nf1* cKO coronal tissue was visualized for tdTomato (*Nkx2.1-Cre*-lineages) and either Olig2 (**A, E, I**), ChAT (**B, F, J**), SST (**C, G, K**) or PV (**D, H, L**). Cell density quantification of tdTomoto+ cells co-labeled for Olig2+ (**M**), ChAT (**N**), SST (**O**) and PV (**P**). Quantification of co-localized tdTomato/SST+ cell density did not reveal differences between genotypes (**E**). All data are expressed as the mean ± SEM. WT, n = 4, cHet, n = 3 and cKO, n = 3. * P < 0.05, **** p < 0.0001. Scale bar in **L** = 100 µm.

**Supplemental Figure 5**

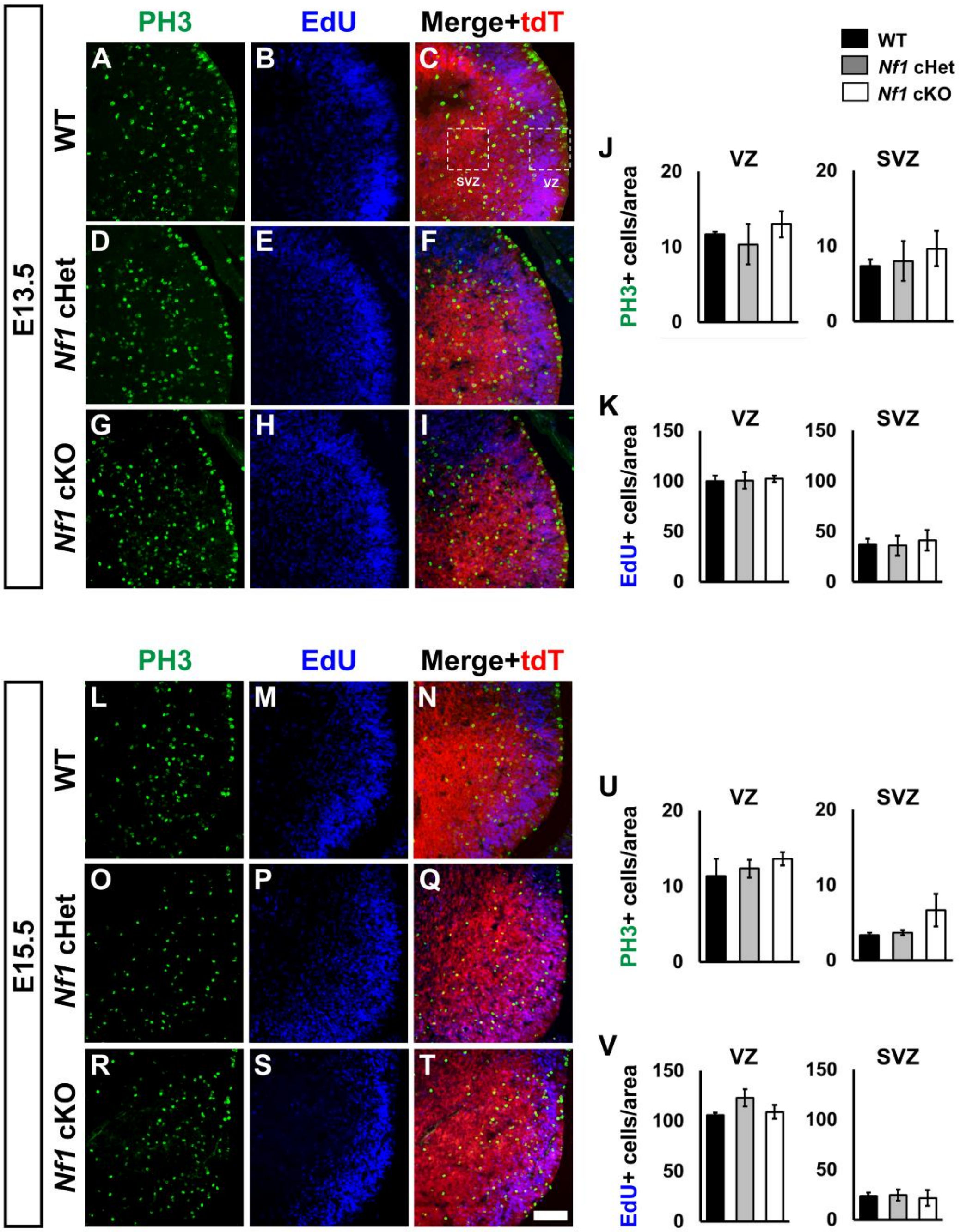


**Supplemental Figure 5: Normal proliferation in the MGE of *Nf1* mutants.**

E13.5 WT (**A-C**), *Nf1* cHet (**D-F**) or *Nf1* cKO (**G-I**) coronal brain sections were co-labeled for PH3, EdU and tdTomato (tdT). Inset boxes in (**C**) denote the ventricular zone (VZ) and sub ventricular zone (SVZ) regions that cells were counted from. Quantification of the cell density of either PH3+ cells (**J**) or EdU+ cells (**K**) in both the VZ and SVZ at E13.5. E15.5 WT (**L-N**), *Nf1* cHet (**O-Q**) or *Nf1* cKO (**R-T**) sections co-labeled for PH3, EdU and tdTomato (tdT). Quantification of the cell density of either PH3+ cells (**U**) or EdU+ cells (**V**) in both the VZ and SVZ at E15.5. Data are expressed as the mean ± SEM. n = 3, all groups. Scale bar in (**T**) = 100 μm.

**Supplemental Figure 6**

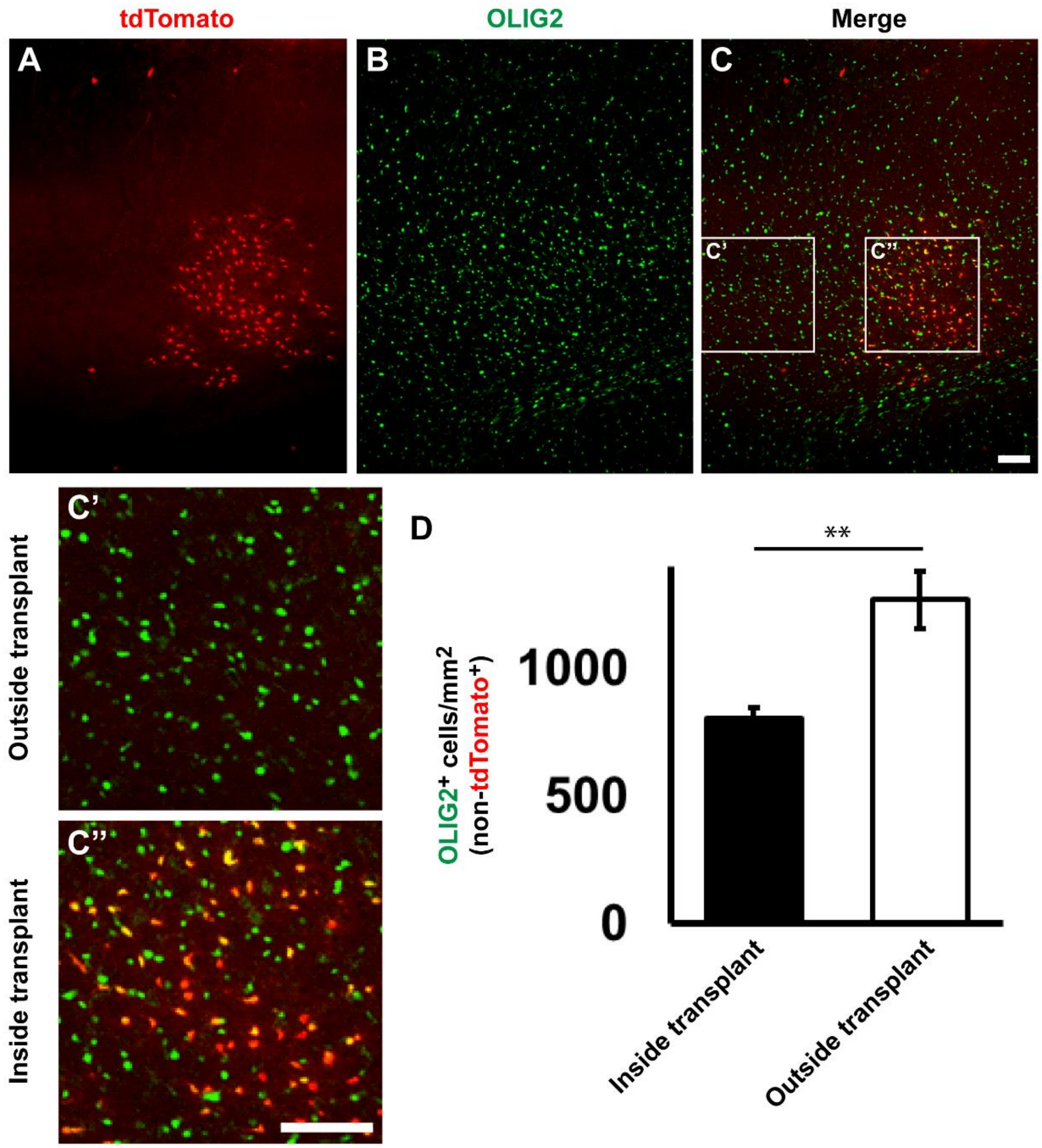


**Supplemental Figure 6: Xenografted *Nf1* cKO oligodendrocytes locally decrease the numbers of WT oligodendrocytes.**

E13.5 cKO MGE cells (tdTomato+) were transplanted into P2 WT cortices and allowed to develop *in vivo* for 35 days. The cortices containing the transplanted cells were co-labeled for tdTomato and OLIG2, revealing discrete clusters of transplanted MGE-derived oligodendrocytes (**A-C**). Boxes were drawn inside the transplant region or outside of the transplant region and OLIG2+ cells that were not tdTomato+ counted in each domain. (**C'** and **C''**) Example higher magnification images of the regions inside and outside of the transplant domain. (**D**) Quantification of non-tdTomato+ cells that are OLIG2+ in each domain. Data are expressed as the mean ± SEM. n = 4 transplant domains and adjacent sites from 2 transplant recipient mice. ** $p < 0.01$. Scale bar in (**C**) = 100 µm.

**Supplemental Figure 7**

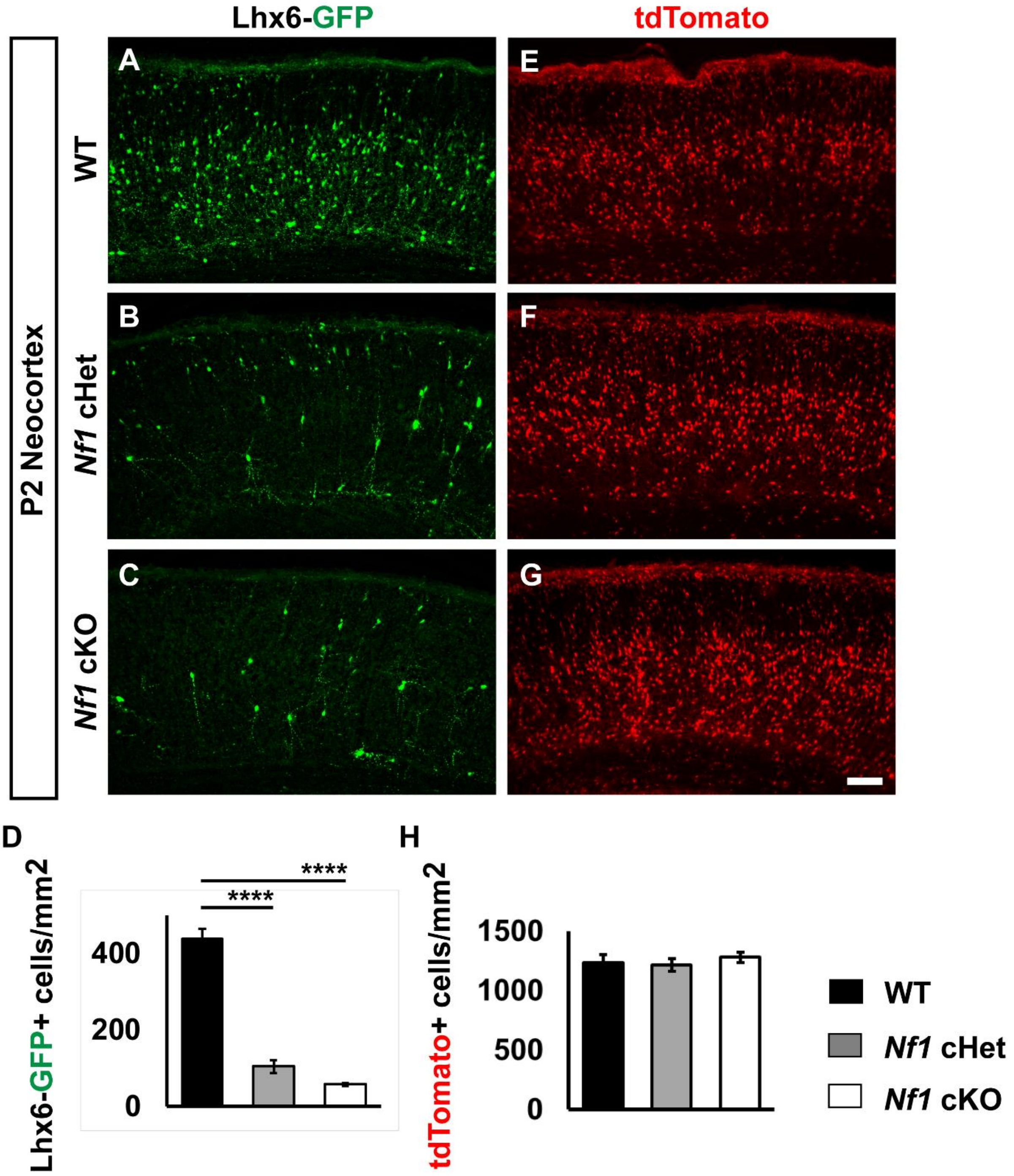


**Supplemental Figure 7*: Lhx6-GFP* numbers are reduced in cHets and cKOs at P2.**

P2 WT, cHet and cKO neocortices were labeled for *Lhx6-GFP* expression (**A-C**). (**D**) Quantification of the cell density of Lhx6-GFP+ cells in the neocortex. P2 brains from the same genotypes were visualized for tdTomato (Nkx2.1-Cre-lineages) (**E-G**) and quantified to determine cell density (**H**). Data are expressed as the mean ± SEM. n = 4, WT/tdTomato counts, n = 3, all other groups. **** $p < 0.0001$. Scale bar in (**G**) = 100 µm.
